# Manipulation of CA1 neuronal subtypes through Cre-mediated viral delivery in mice

**DOI:** 10.64898/2026.05.08.723440

**Authors:** Dheeraj Songara, Hiyaa S. Ghosh

## Abstract

CaMKIIα promoter is widely used to label and manipulate hippocampal pyramidal neurons via transgenic mouse lines or viral approaches. While it targets most excitatory neurons, a small subset remains unlabeled and often overlooked. We present an AAV-based strategy combined with CaMKIIα-driven Cre expression to access and study this remaining population. Furthermore, we provide a detailed protocol for in-house AAV production, targeted stereotaxic delivery, and functional validation of targeted neurons through slice electrophysiology and behavior.

**Graphical abstract:** 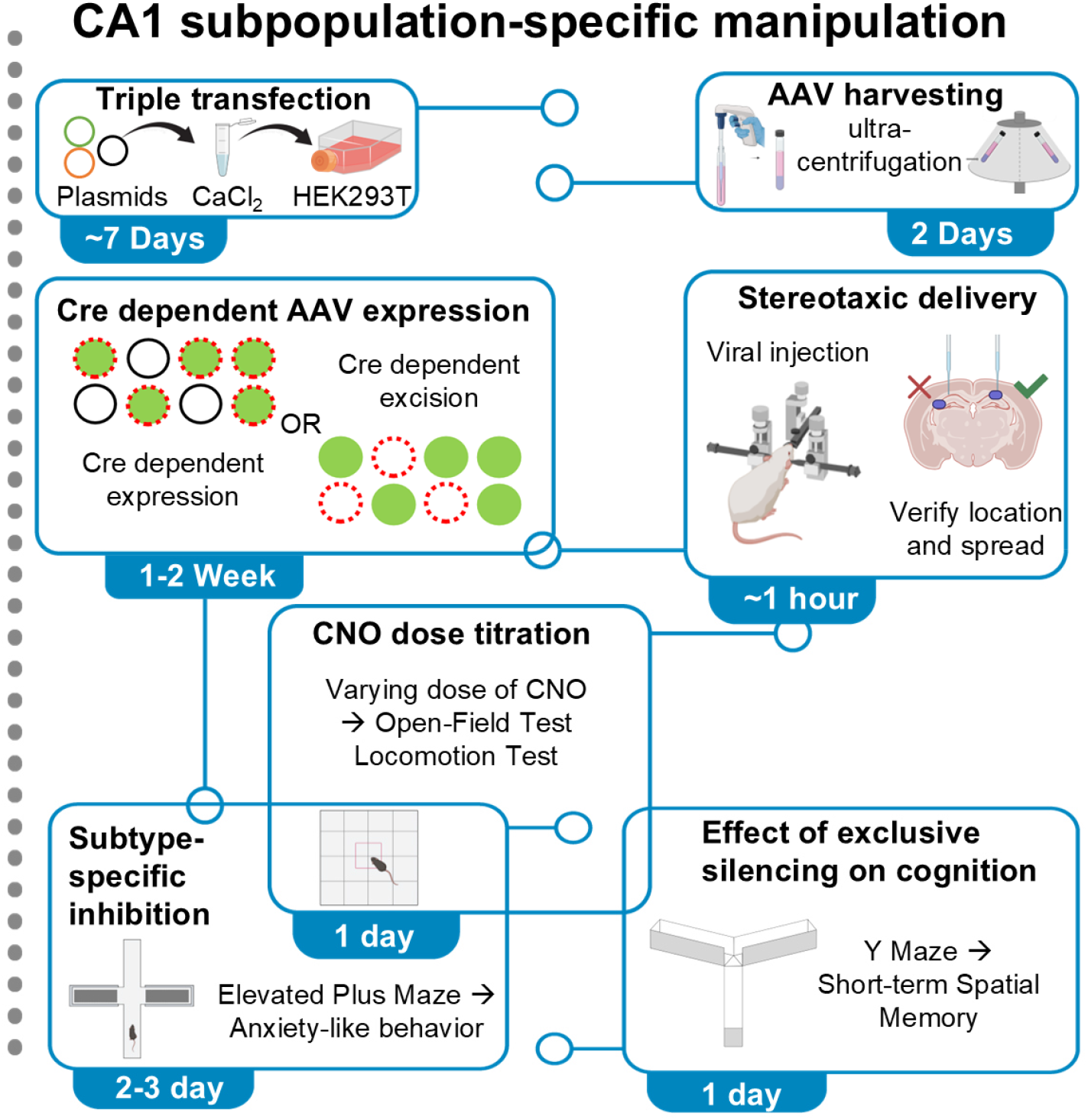

## Introduction

This protocol describes the specific steps for AAV production and stereotaxic delivery into the dorsal CA1 of adult CaMKIIα-CreERT2 mice to differentially target Cre-positive (CaMKIIα+) and Cre-negative (CaMKIIα-) excitatory neuronal subpopulations. The same approach can be adapted for other brain regions by modifying the stereotaxic coordinates accordingly ^1^.

The CaMKIIα promoter is widely used to target excitatory pyramidal neurons in the hippocampus, and its use for labeling excitatory neurons has been well established in both AAV-based and transgenic approaches^2–5^. Critically, CaMKIIα-mediated approaches have been critical in establishing a role for CA1 neurons in spatial and fear memory in studies related to learning and memory^6–8^. However, previous work by our group and others show that CaMKIIα-driven approach only captures ∼70% CA1 neurons^9,10^. This leaves an important caveat of excluding the remaining ∼30% CA1 neurons in studies that use CaMKIIα-driven approaches in their experiments. In order to address this caveat, we developed a strategy that uses a careful combination of AAV plasmids to use a Cre-dependent ON/OFF system to target both CaMKIIα+ as well as CaMKIIα-CA1 neurons using CaMKIIα-Cre AAV mediated DREADD delivery.

Furthermore, gene delivery through viral vectors, and adeno-associated viruses (AAV) in particular, has become an indispensable tool across many fields of biomedical research^11–13^. Unlike transgenic animal models, which are expensive to generate and maintain, require months to achieve the desired genotype, and offer limited spatial control over gene expression, AAV-based approaches allow researchers to target specific cell populations in specific anatomical regions with remarkable precision^14,15^. By combining AAV serotypes with carefully engineered plasmid designs incorporating cell-type-specific promoters, Cre- or Flp-dependent recombination switches, and intersectional strategies, investigators can achieve a level of experimental control that transgenic lines simply cannot offer^16–18^ This flexibility, combined with the broad tissue tropism of naturally occurring and engineered AAV capsids^19,20^, has driven the widespread adoption of all-viral approaches across neuroscience, gene therapy, and beyond.

Despite this enthusiasm, access to high-quality AAV remains a persistent bottleneck for many research groups. Commercial vendors can supply well-characterized, ready-to-use preparations, but the associated costs are prohibitive, particularly for laboratories requiring multiple constructs or large volumes. Beyond cost, international shipping on dry ice, customs clearance delays that can stretch into months, and the logistical fragility of the cold chain make vendor-sourced AAV an unreliable option for many institutions. These barriers have motivated a growing number of laboratories to bring AAV production in-house^21,22^.

However, in-house production carries its own set of challenges. The available literature describes a wide variety of upstream transfection strategies-including transient triple transfection in HEK293 or HEK293T cells^23–25^, as well as numerous downstream purification approaches ranging from iodixanol density gradient ultracentrifugation^21,26^, to affinity and ion-exchange chromatography^27,28^. While each of these methods has been validated in specialized production facilities, translating them into a standard academic laboratory setting is far from straightforward. Many protocols rely on expensive proprietary kits and reagents, require specialized equipment, or involve multi-day, labor-intensive workflows that are difficult to standardize without prior expertise. In our own experience, arriving at a reliable, reproducible production pipeline took considerable time and iteration before we could trust the quality and consistency of the resulting vector preparations.

Here, we describe a streamlined AAV production protocol built around widely available, low-cost reagents and standard laboratory equipment. Our upstream workflow uses calcium-phosphate transfection of HEK293T cells with a three-plasmid system, avoiding the need for commercial transfection reagents. Purification is achieved through sucrose density gradient ultracentrifugation - steps that can be completed within one week, from cell seeding to a ready-to-use vector preparation. We also describe our approach for quality control, including titer determination by qPCR and functional validation by in vitro transduction. The goal of this paper is not to introduce a fundamentally new method, but to consolidate existing knowledge into a practical, accessible, and fully reproducible guide that any laboratory, regardless of prior experience, can implement without prohibitive cost or infrastructure.

Finally, to validate the targeting of CaMKIIα-Cre driven approach for both Cre-positive and Cre-negative excitatory neuronal subpopulations, we use a short-term spatial learning and memory in mice expressing hM4Di in dorsal CA1.

## Materials

| REAGENT or RESOURCE | SOURCE | IDENTIFIER |
| --- | --- | --- |
| **Bacterial and virus strains** |  |  |
| pHelper (pAdDeltaF6) | James M. Wilson | Cat# 112867; RRID: Addgene_112867 |
| pAAV2/1 (RepCap) | James M. Wilson | Cat# 112862; RRID: Addgene_112862 |
| pAAV-hSyn-FLEX-Loxp-hM4di-mCherry | Beier et al. | Cat# 161581; RRID: Addgene_161581 |
| pAAV-hSyn-FLEX-OFF-hM4di-mCherry | Beier et al. | Cat# 161579; RRID: Addgene_161579 |
| **Chemicals; peptides; and recombinant proteins** |  |  |
| Tamoxifen | TRC | Cat# T006000 |
| Clozapine-N-oxide | HelloBio | Cat# HB6149 |
| Paraformaldehyde | Himedia | Cat# GRM3660 |
| Anti-Anti (Penicillin–Streptomycin–Amphotericin B) | Gibco; Thermo Fisher Scientific | Cat# 15240062 |
| Opti-MEM™ I Reduced Serum Medium | Gibco; Thermo Fisher Scientific | Cat# 31985062 |
| Fetal Bovine Serum | Gibco; Thermo Fisher Scientific | Cat# 16140071 |
| DMEM, high glucose, GlutaMAX™ Supplement | Gibco; Thermo Fisher Scientific | Cat# 10566016 |
| TrypLE | Gibco; Thermo Fisher Scientific | Cat# 12604013 |
| DNase I | Sigma-Aldrich | Cat# D4513 |
| Clozapine-N-oxide (CNO) | Hellobio | Cat# HB6149 |
| **Experimental models: Cell lines** |  |  |
| AAVpro 293T | Takara Bio | Cat# 632273; RRID: CVCL_BOXW |
| **Experimental models: Organisms/strains** |  |  |
| Mouse: B6;129S6-Tg (Camk2a-cre/ERT2)1Aibs/J | The Jackson Laboratory | Strain#:012362; RRID: IMSR_JAX:012362 |
| Mouse: STOCK Gt (ROSA)26Sortm1.1(CAG-EGFP) Fsh/Mmjax | The Jackson Laboratory | Strain#032037; RRID: MMRRC_032037-JAX |
| **Software and algorithms** |  |  |
| ImageJ/Fiji v1.54p | NIH | RRID: SCR_002285 |
| ANY-maze v7.4 | Stoelting | RRID: SCR_014289 |
| pClamp / Clampfit | Molecular devices | RRID:SCR_011323 |
| **Other (Instruments)** |  |  |
| Micropipette puller | Sutter Instrument | P-1000 |
| Vibratome | Leica Microsystems | VT1200S |
| Behavioral tracking camera | Custom-built | N/A |
| PicoSpritzer | Parker Hannifin | PicoSpritzer III |
| Stereotaxic frame | Kopf Instruments | Model 940 |

## Equipment

- Biosafety cabinet, Class II
- CO_2_ incubator (37°C, 5% CO_2_)
- Benchtop centrifuge
- Ultracentrifuge or high-speed centrifuge (≥23,000 × g)
- Water bath or bead bath
- -80°C freezer
- Hemocytometer
- Epifluorescence microscope (inverted)
- Confocal microscope (optional)
- qPCR machine
- Stereotaxic setup
- Stereoscope
- Isoflurane delivery chamber and nose cone
- PicoSpritzer III and stimulator
- Pipette puller
- Heating pad and temperature controller
- Vibratome
- Carbogen (95% O_2_ / 5% CO_2_) supply
- Nitrogen supply for PicoSpritzer

### Reagents

**2X HEPES-Buffered Saline (HBS)** (as per Cold Spring Harbor Laboratory (CSHL) guidelines)

| Reagent (stock) | Final concentration | Amount |
| --- | --- | --- |
| D-Glucose (100 mg/mL) | 10 mM | 0.18 mL |
| HEPES (1 M) | 40 mM | 0.40 mL |
| KCl (1 M) | 10 mM | 0.1 mL |
| NaCl (5 M) | 270 mM | 0.54 mL |
| Na<sub>2</sub>HPO<sub>4</sub> (0.5 M) | 1.5 mM | 0.030 mL |
| ddH<sub>2</sub>O | n/a | Up to 10 mL |
| **Total** | **n/a** | **10 mL** |

Adjust pH to 7.1 with NaOH after mixing all components. Bring final volume to 10 mL with ddH2O. Filter-sterilize and aliquot into 250 μL. It can be stored at -20°C up to several months.

**Alternatives**: 2X HBS can be bought commercially.

#### Plasmid mix

| Reagent | Final concentration | Amount |
| --- | --- | --- |
| RepCap plasmid | n/a | 10 µg |
| Helper plasmid | n/a | 20 µg |
| Transfer plasmid | n/a | 10 µg |
| CaCl<sub>2</sub> (2.5 M) | 250 mM (final approx.) | 25 µL |
| ddH<sub>2</sub>O / NFW | n/a | Up to 250 µL |
| **Total** | **n/a** | **250 µL** |

**Note:** Calculate the amount of water and add the water first. Prepare 5 minutes before use.

**CRITICAL**: Ensure sterile conditions while preparing the plasmid mix. CaCl2 is hygroscopic; prepare fresh or use properly stored stock. Avoid vortexing DNA to prevent shearing.

**Alternatives**: Commercial transfection reagents (e.g., PEI, Lipofectamine) can be used instead of calcium phosphate method depending on application.

#### Complete DMEM culture media

| Reagent | Final concentration | Amount |
| --- | --- | --- |
| DMEM | n/a | Appropriate volume |
| Fetal Bovine Serum (FBS) | 10% (v/v) | 5 mL per 50 mL media |
| Anti-Anti (100X) | 1X | 0.5 mL per 50 mL media |
| **Total** | **n/a** | **50 mL** |

Filter-sterilize using a 0.22 μm filter and store at 4°C. Warm to 37°C before use.

**CRITICAL**: Use sterile technique while preparing media to avoid contamination. Do not repeatedly freeze-thaw FBS.

**Alternatives**: Penicillin-Streptomycin can be used instead of Anti-Anti depending on experimental requirements.

**20% Sucrose Solution (Tris-based buffer)**

| Reagent | Final concentration | Amount (for 100 mL) |
| --- | --- | --- |
| Tris HCL | 50 mM | 5 mL of 1 M Tris pH 7.4 |
| NaCl | 100 mM | 0.584 g |
| EDTA | 0.5 mM | 100 µL of 0.5 M EDTA |
| Sucrose | 20% (w/v) | 20 g |
| ddH<sub>2</sub>O | n/a | Up to 100 mL |
| **Total** | **n/a** | **100 mL** |

Adjust pH to 7.4 using HCl, filter-sterilize (0.22 μm), and store at 4°C for several months.

**CRITICAL**: EDTA may take time to dissolve; adjust pH slightly to aid dissolution.

#### 4% Paraformaldehyde (PFA)

| Reagent | Final concentration | Amount |
| --- | --- | --- |
| Paraformaldehyde | 4% | 4 g |
| 10X PBS | 1X PBS | 10 ml |
| Milli-Q | n/a | 90 ml |
| **Total** | **n/a** | **100 mL** |

Pre-heat the 1X PBS to 60°C in a microwave and keep it on a heating stirrer to maintain. Add the PFA powder and let it dissolve for 15 minutes.

**CRITICAL**: PFA is a toxic and biohazardous chemical. Wear proper PPE before dealing with PFA and work under a fume hood. Store at 4°C up to a week.

## Procedure

Experiments were conducted on both male and female mice in accordance with the protocol approved by the Institutional Animal Care and Ethics Committee (IACEC). The mice used in the study were bred and housed in the Institutional Animal Care and Resource Center (ACRC) in SPF barrier facility. The animals were housed in their regular individually ventilated cages with easy access to food and water with 14/10 hours of light/dark cycle.

### Cre-dependent (Flex-OFF) Viral Strategy

To selectively target and label Cre-negative excitatory neurons, we used a Cre-dependent “Flex-OFF” AAV vector. In this design, the hM4Di-mCherry coding sequence is placed in the correct orientation for transcription and flanked by heterotypic loxP and lox2272 sites^29^. In the absence of Cre recombinase (Cre-neurons), the cassette remains intact and drives robust expression under the human Synapsin (hSyn) promoter. In the presence of Cre (Cre+ neurons), recombination between lox sites causes irreversible excision of the coding sequence, thereby preventing expression. This strategy ensures that only Cre excitatory neurons are labeled. The virus was stereotaxically injected into the dorsal CA1 region of CaMKIIα-CreERT2 mice, and animals were allowed 3-4 weeks for expression before downstream validation.

### AAV production via Calcium-Phosphate Transfection

**CRITICAL**: All the calculations are done for T25 flask, in order to increase the viral quantity, it is recommended to put multiple T25 flasks.

#### 1. Cell Preparation: Timing: [<1 week]

**Note:** Use HEK cells with fewer than 10 passages for optimal viral production. AAVpro 293T cells, a variant of HEK293T, are recommended for higher AAV yield.

a. Prepare culture media by adding 10% Fetal Bovine Serum (FBS) and 1x Anti-Anti to DMEM, sterilize the media using a 0.22 μm filter and store at 4°C.
b. Warm the media to 37°C before starting the culture.
c. After reviving cells, passage them a few times to achieve optimal growth rate.
d. Split the cells one day before transfection to achieve 70% confluency the next day.
e. Split the cells by removing the media and wash the cells with 1 mL of fresh media to remove floating cells and debris.
f. Add 1 mL of TrypLE directly to the cells and incubate at 37°C for 3 minutes.
g. Gently tap the flask to dislodge cells, and collect them at one corner of the flask.
h. Triturate the detached cells several times using a 1 mL pipette.
i. Add 2 mL of fresh media to dilute TrypLE and collect it 15 ml falcon tube.
j. Centrifuge the cell suspension at 250 x g for 5 minutes at room temperature
k. Discard the supernatant and resuspend the cell pellet in 1 mL of fresh media. Ensure the cells are evenly dispersed.

**CRITICAL:** Avoid clumps, as they can interfere with transfection efficiency.

l. Seed 1–3 million cells in a T25 flask, depending on the growth rate of your cell line.

**CRITICAL:** If the cells are freshly revived from frozen stocks, split them at least twice before seeding to optimize their condition.

m. To determine the best cell density, seed multiple flasks with varying cell numbers.
n. Aim for ∼70% confluency the following day for transfection.
o. Avoid overgrowth beyond 24 hours before transfection.

#### 2. Transfection Protocol: Timing: [1 hour]

a. Replace the culture media in the T25 flask with pre-warm Opti-MEM at least 1 hour before transfection.
b. Thaw a 250 μL of 2x HBS for use in a 5 mL tube.
c. Mix the plasmid solution thoroughly by pipetting several times.
d. Take a drop of plasmid mix solution with 1 ml pipette and add it to the 2x HBS kept in 5 ml tube to make calcium phosphate-plasmid precipitate.

**Note:** Ensure slight bubbling during mixing; aeration improves the formation of calcium phosphate-DNA precipitates.

e. Repeat the dropwise addition until the entire solution is mixed.
f. Immediately add the prepared calcium phosphate-plasmid precipitate solution (within 5 minutes of preparation) to the T25 flask containing cells in Opti-MEM.
g. Evenly distribute the solution by carefully dropping it over all regions of the flask.
h. Gently swirl the flask to ensure uniform coverage.
i. Incubate the cells at 37°C in a 5% CO_2_ atmosphere within a BSL-2 facility.
j. The following day, carefully replace the media with FBS-free DMEM.
k. Check fluorescence intensity and cell health daily using an inverted epifluorescence microscope.

**Note:** Confirm fluorescence before proceeding with media collection. If the fluorescence is absent in your plasmid, perform a parallel transfection with a fluorescent marker using the same reagents to ensure transfection success.

l. Upon seeing > 30-40 % fluorescence labelled cells, collect media after 48- and 72-hours post-transfection and store at 4°C.

**Note:** Cell heath may deteriorate after transfection but if you see good fluorescence then you can still harvest the AAV.

m. If cells remain healthy, collect additional media at 96 hours.

**CRITICAL**: Do not extend collection beyond 96 hours to prevent viral particle degradation.

#### 3. Harvesting: Timing: [2 days]

a. Pool all the harvested media into a 15 ml Falcon tube.
b. Centrifuge the media at 2000 rpm for 10 minutes at 4°C to remove debris.
c. Collect the supernatant and filter it using a 0.22 μm filter directly into 10 ml ultracentrifuge tubes.

**Note:** If you have more than 6 ml of media, use two ultracentrifuge tubes. Ultracentrifuge tubes can be reused after proper washing with bleach and soap, followed by thorough drying.

d. Slowly add the 20% sucrose solution in a 1:4 ratio directly to the bottom of the ultracentrifuge tube.
e. Mark the direction of pellet formation on the cap of the ultracentrifuge tube and centrifuge at 23000 × g for 3 hours at 4°C.
f. Carefully remove the tubes and circle the location of pellet formation.
g. Remove the supernatant, tilt the tube so that the pellet remains on the upper side, let it sit for 2-3 minutes, and remove any excess supernatant.
h. To dissolve the pellet, add 50 μl of chilled 1× PBS directly to the pellet, keeping the tube tilted so that the pellet is submerged. Let it dissolve overnight at 4°C.
i. The next day, resuspend the pellet slowly using a 20 μl pipette multiple times without introducing bubbles. Pool the virus from all the tubes and mix thoroughly.
j. Aliquot into 5 μl volumes and store at -80°C.

#### 4. qPCR: Timing: [3 hours]

a. Quantify viral genome per μL (vg/μL) using qPCR against a plasmid standard curve following the Addgene protocol (<u>link</u>).

### Stereotaxic Injection

#### Timing: [1-2 hour per animal for bilateral injection]

**CRITICAL:** Perform the viral injections in a class II biosafety hood.

#### 5. Injection setup

a. Turn on the heating pad before starting the experiment and open the carbogen and nitrogen supply valves.
b. Adjust the carbogen supply pressure in the isoflurane chamber to ∼0.9 and set the nitrogen pressure in the main line valve to 20 pascal and ∼15-20 pascal for the PicoSpritzer.
c. Use a pedal remote to deliver air pulses of <10 ms.
d. Anesthetize the mouse and place it on the stereotaxic setup. Adjust the isoflurane level to 2-3%, depending on the mouse’s breathing reflex (target: 0.5–1 Hz).
e. Visually align and position the mouse’s head straight.
f. Apply eye wax to prevent the eyes from drying.
g. Decontaminate the head by applying betadine with a sterile cotton swab.

#### 6. Calibration of Injection Parameters: Timing: [1-2 weeks]

Before commencing for viral injection, calibrate the injection speed and volume using a diluted tracer dye. This step is essential for ensuring accurate targeting and predictable viral spread, and does not require virally-injected animals.

##### a. Injection Speed and Spread Calibration

I. Dilute trypan blue 1:10 in sterile 1× PBS.
II. Inject 1 μL of diluted dye at three different infusion rates (e.g., 0.05, 0.10, and 0.20 μL/min) into the dorsal CA1 of separate animals.
III. Immediately sacrifice each animal after injection and extract the brain into 1× PBS.
IV. Section the brain manually or with a vibratome into 300-500 μm slabs through the injection site to visualize dye spread and confirm targeting (Figure 4C).

**Figure 1:**
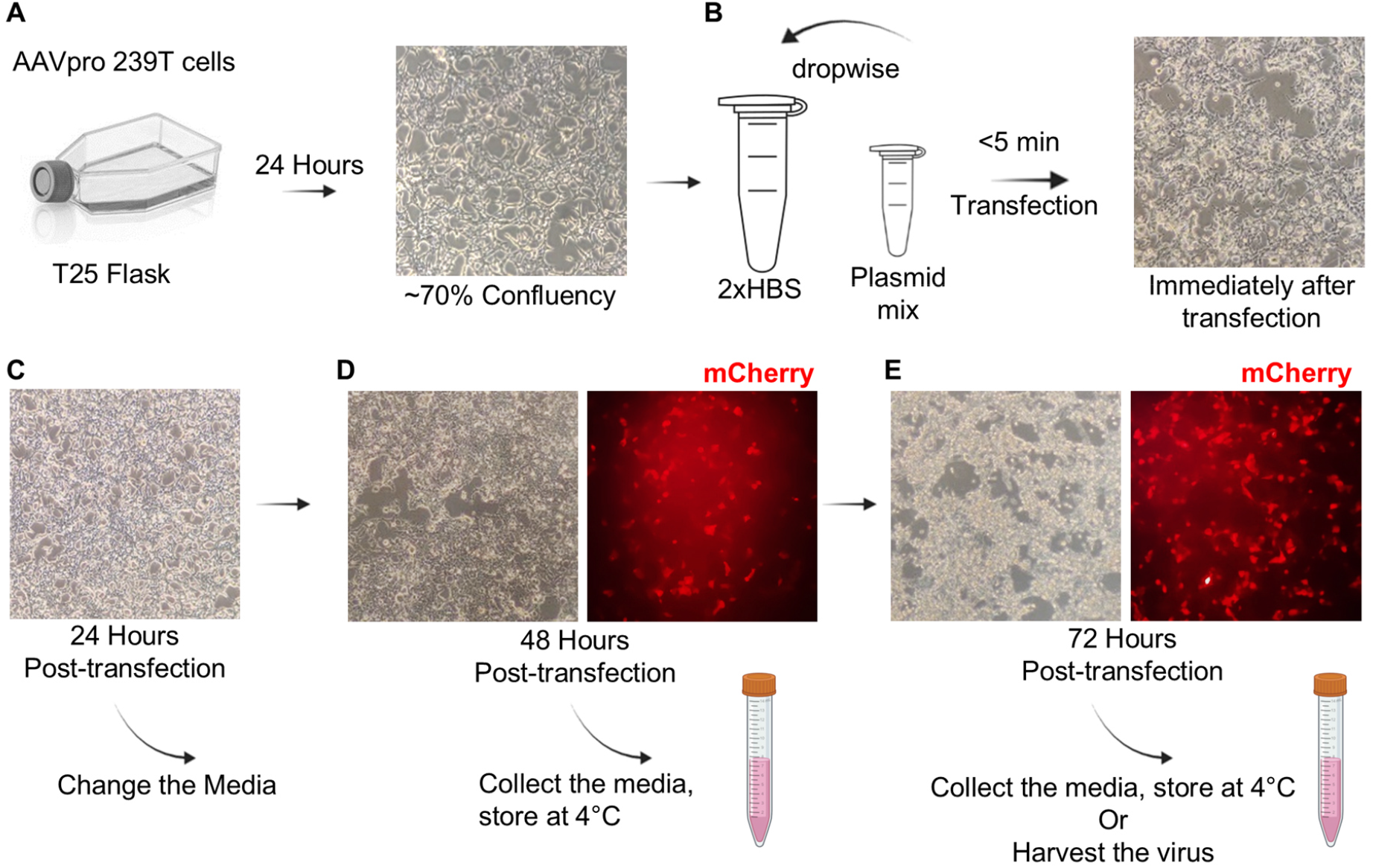
AAV Production via Triple Transfection in HEK293T Cells. (A) AAVpro 293T cells are seeded at a density of 3.0–3.5 million cells in a T25 flask and allowed to reach ∼70% confluency in 24 hours. (B) Triple transfection is performed using 2.5 M CaCl_2_ with 2xHBS buffer and three plasmids: a transfer plasmid, a helper plasmid, and a RepCap plasmid. (C) Media change is done 24 hours after transfection. (D-E) Transfection success is monitored via reporter expression (e.g., mCherry), visible at 48–72 hours. Media is collected at 48- and 72-hours post-transfection and stored at 4°C. The virus-containing media is later used for purification.

**Figure 2.**
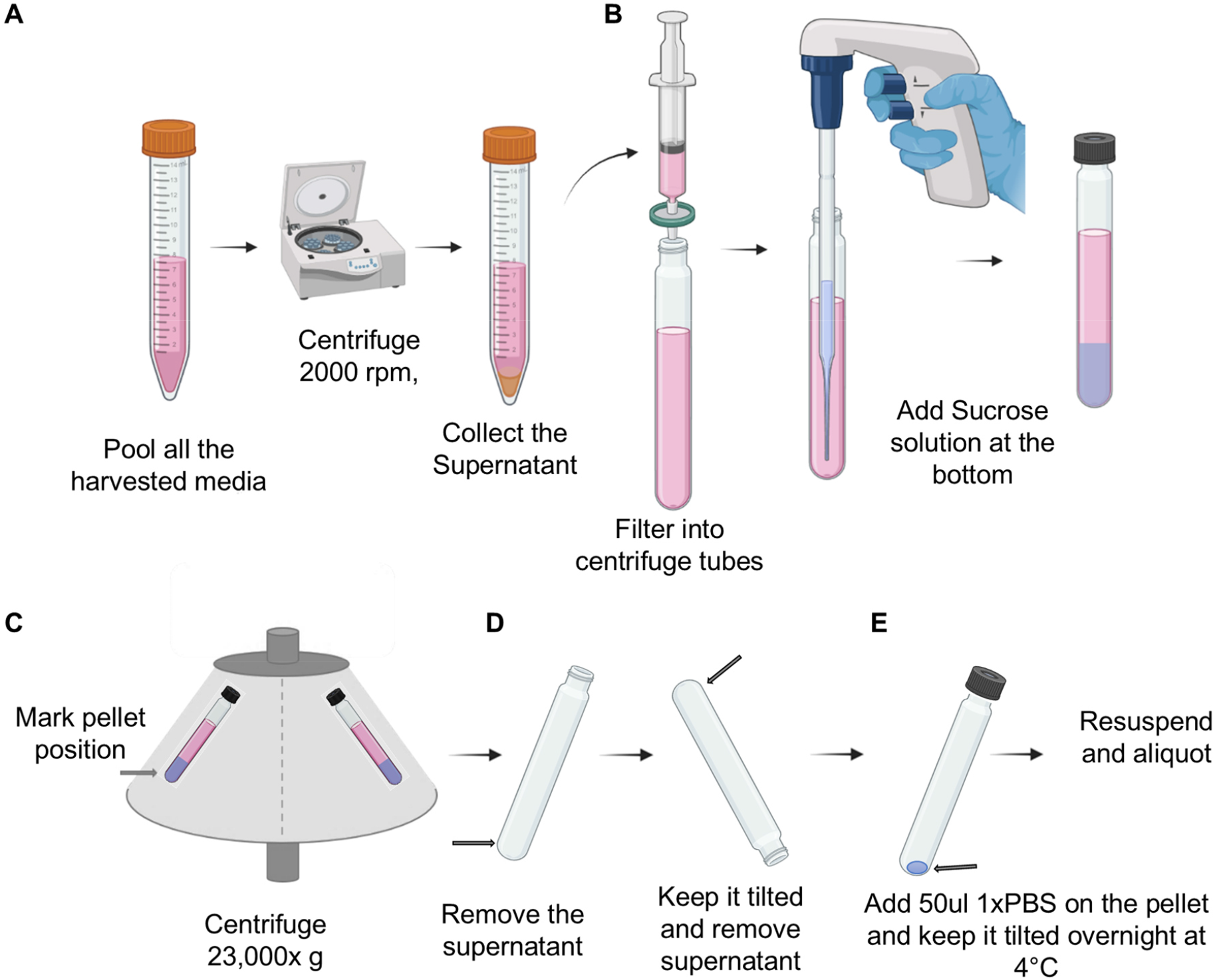
AAV Purification via Sucrose Cushion Ultracentrifugation. (A) Harvested viral media is pooled and centrifuged at 2000 rpm for 10 minutes at 4°C to remove cellular debris. (B) The supernatant is filtered through a 0.22 μm filter and underlaid with a 1:4 volume ratio of 20% sucrose solution. (C) The mixture is ultracentrifuged at 23,000 ×g for 3 hours at 4°C. (D) The supernatant is carefully removed, and the tube is tilted to drain residual fluid. (E) The viral pellet is resuspended in 50 μL of 1× PBS and incubated overnight at 4°C. Final resuspension is done using a 20 μL pipette, make 5 μL aliquots and storage in -80°C. Arrow indicates the location of expected pellet.

**Figure 3.**
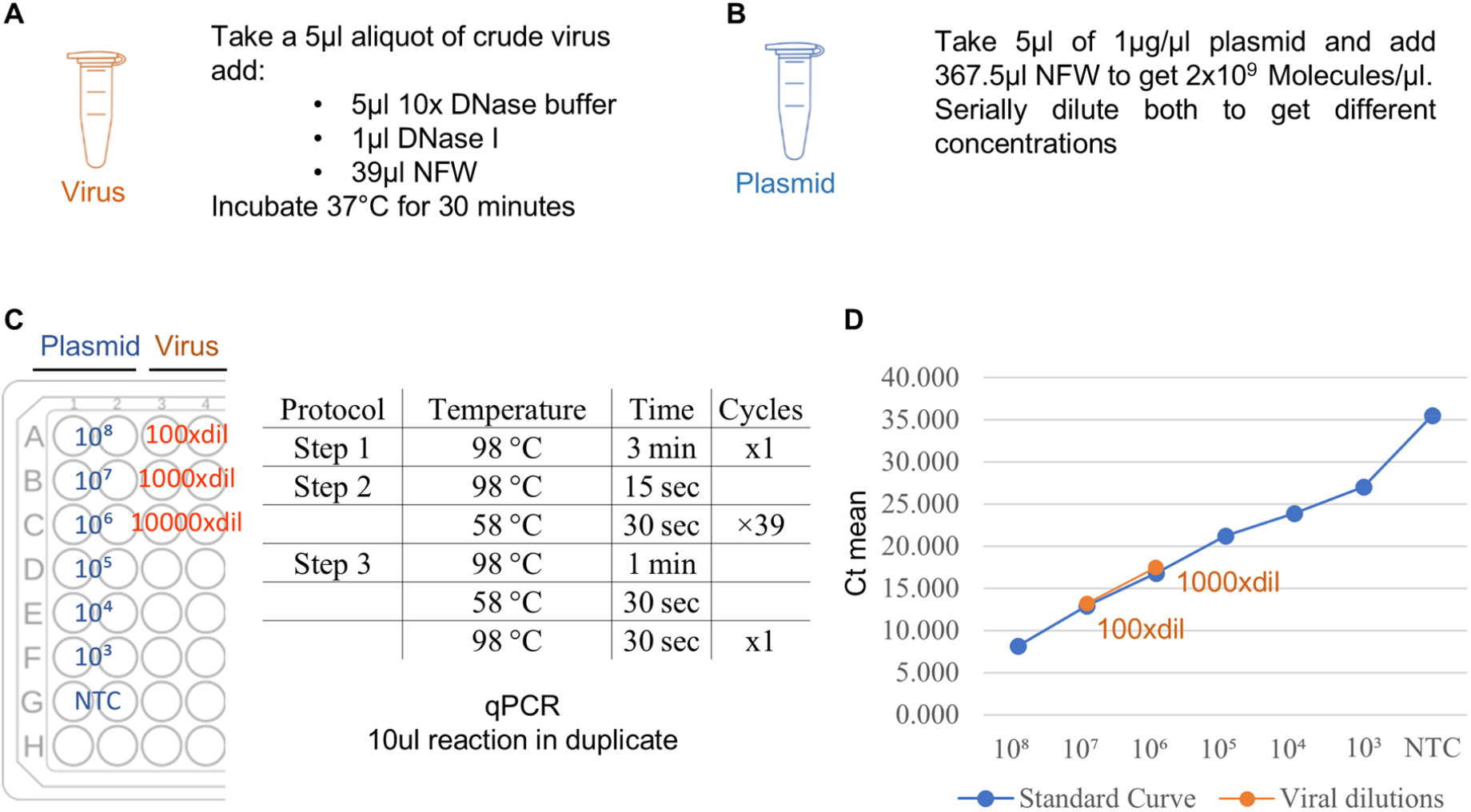
Viral Titer Determination via Quantitative PCR. (A) A 5 μL aliquot of crude virus is treated with DNase I to eliminate residual plasmid DNA, followed by incubation at 37°C for 30 minutes. (B) A standard curve is generated using serial dilutions of a plasmid of known concentration. (C) qPCR is run with duplicate 10 μL reactions using a denaturation step at 98°C followed by 39 cycles of amplification and a melt curve step. (D) Viral samples are diluted (e.g., 1:100 and 1:1000) and compared to the plasmid standard curve to estimate viral genome copies.

**Figure 4.**
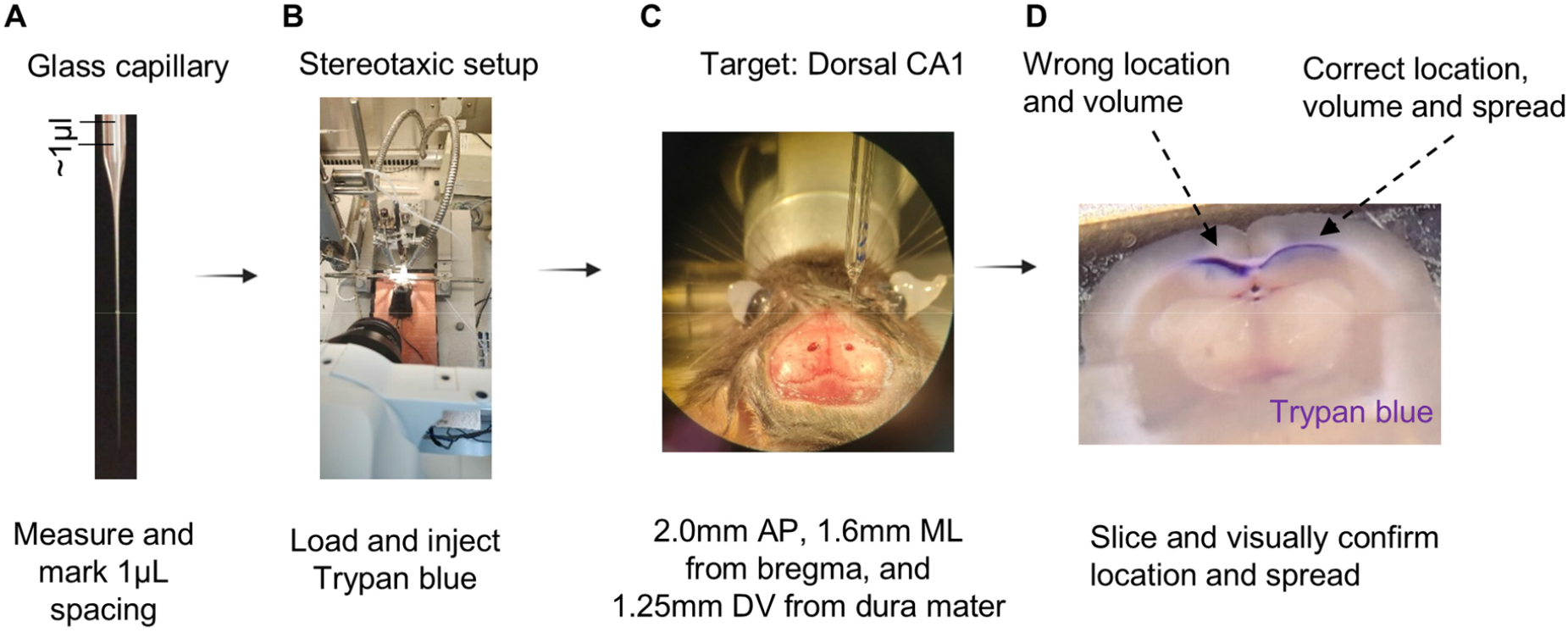
Stereotaxic Injection of AAV into the Mouse Dorsal CA1 Region. (A) Stereotaxic injection of AAV is performed into the dorsal hippocampus using glass capillary. (B) Trypan blue is used as a test dye to optimize injection parameters, including location, speed, and volume. (C) Injection coordinates are AP: -2.0 mm, ML: ±1.6 mm from bregma, and DV: -1.25 mm from the dura were used to target dorsal CA1. (D) Post-injection brains were sliced to confirm correct targeting and diffusion by visual inspection of Trypan blue spread. Correct injections show clear localization within the dorsal CA1 region.

**Note**: Select the injection rate that produces the most contained, spherical spread at your target volume. Overly rapid injection can cause reflux along the needle tract.

##### b. Viral Volume and Titer Optimization

i. Inject varying volumes (e.g., 0.3, 0.5, and 1.0 μL) of virus into separate animals at the optimized injection rate.
ii. Sacrifice animals 5-7 days post-injection and perfuse transcardially with 1× PBS followed by 4% PFA.
iii. Section brain tissue at 30-50 μm around the injection site. Stain sections with DAPI and evaluate fluorescence coverage using epifluorescence or confocal microscopy.

**Note**: Select the AAV volume that achieves adequate fluorescence coverage of the dorsal CA1 with minimal extravasation to adjacent regions.

#### 7. Stereotaxic Viral Injection: Timing: [1-2 hours per animal]

a. Using forceps, lift the skin from mid line near the ear position and cut a small section to expose the skull, ensuring the bregma and lambda are visible for alignment.
b. Clean and dry the skull, removing connective tissue with a fresh cotton swab.
c. Fill the glass pipette with virus, ensuring no air bubbles. Mark the volume levels on the pipette using a marker.
d. Attach the glass pipette to the pipette holder. Cut the pipette tip slightly, and apply air pulses until a small drop of virus forms and stays at the pipette tip. If it does not stay, cut the tip a bit more.

**CRITICAL**: Air bubbles within the capillary will cause inconsistent injection volumes and should be removed before proceeding.

e. To align the mouse head correctly, adjust the bregma and lambda to the same level (±0.05 mm). Adjust the lateral tilt by checking the coordinates by moving 1 mm left and right from the bregma-lambda midpoint and ensuring the same levels.
f. Move to the desired coordinate and drill the skull until a thin layer remains.
g. Use a 26-gauge needle to gently create a hole by scraping the skull. If bleeding occurs, clean it until it stops.
h. Lower the pipette while applying air pressure to prevent blockages. Mark the surface as zero for the dorsoventral coordinate. While apply air pressure slowly lower the pipette at 0.2 mm/s until reaching the desired depth.
i. Inject the virus using a stimulator to stimulate the PicoSpritzer at 0.3-0.4 Hz, ensuring no blockade by zooming in on the pipette.
j. After completing the injection, wait 2 minutes, then slowly retrieve the pipette.
k. Apply the bone wax in the drilled region. Add the dental cement on the skull covering the entire open region and using a syringe drop wise add the cold cure.
l. Let the mouse recover for 2-3 weeks for the experiments or else just to check the expression, it can be sacrificed with a week.

**Note**: Coordinates for dorsal CA1 in adult C57BL/6 mice: AP -2.0 mm, ML ±1.6 mm from bregma, DV - 1.25 mm from dura (adjust as needed for specific strains and experimental requirements).

### Tissue Fixation and Sectioning

#### 8. Perfusion and brain fixation: Timing: [15 minutes per animal]

a. Deeply anesthetize the mouse and perform transcardial perfusion with ice-cold 1× PBS followed by 4% paraformaldehyde (PFA).
b. Dissect the brain carefully and post-fix overnight at 4°C in 4% PFA.

#### 9. Sectioning: Timing: [1 hour per animal]

a. Section the brain coronally using a vibratome (Leica VT1200 S) to obtain 50 μm thick slices.

### AAV Expression Validation by Microscopy

#### 10. Microscopy: Timing: [1 hour per animal]

a. Select sections containing injected site and perform DAPI staining for nuclear visualization.
b. Mount the stained sections onto slides using appropriate mounting medium.
c. Acquire images using a confocal microscope at 40× magnification.
d. Verify expression of viral-driven fluorescence in different cell types.

### Functional Validation of hM4Di by Ex Vivo Electrophysiology

#### 11. Whole-cell patch-clamp: Timing: [1 day]

Prepare all solutions and acute hippocampal slices using the optimized N-Methyl-D-glucamine (NMDG) protective recovery method as described in Allen Institute for Brain Sciences protocol (<u>link</u>).

a. Briefly, deeply anesthetize the mouse with isoflurane and perform transcardial perfusion using ice-cold NMDG-aCSF.
b. Prepare 250 μm thick coronal brain slices containing dorsal CA1 using a vibratome.
c. Transfer slices to holding-aCSF at 32°C, then maintain at room temperature until recording.
d. Perform whole-cell patch-clamp recordings from visually identified mCherry^+^ CA1 pyramidal neurons using DIC and epifluorescence optics.
e. Use recording pipettes (3–5 MΩ) filled with internal solution.
f. Record baseline membrane potential and firing properties prior to drug application.
g. In case of DREADD receptor hM4Di, bath-apply Clozapine-N-oxide (CNO, 10 μM) for 10 minutes to activate hM4Di receptors.
h. Assess functional activity by measuring hyperpolarization of the resting membrane potential.
i. Confirm suppression of neuronal excitability by absence of action potential firing at 125 pA current injection white holding the cells are -65 mV for CA1 neurons.

### Functional Validation of Cre-Strategy by Behavioral Tests

Open-Field Test (OFT): Locomotion test

Elevated Plus Maze (EPM): Anxiety-like behavior test

Y-maze: Short-term spatial memory test

#### Timing: [1 day per behavior]

a. Handle animals for a minimum of 3 days in the behavior room prior to the start of experiments.
b. Conducted behavioral tests on (same animals) separate days with a 1-2 days interval in the following order: Open Field Test (OFT), Elevated Plus Maze (EPM), Y-Maze.
c. Acclimatize animals to the behavior room for at least 10-15 minutes before each test.
d. Clean the apparatus with 70% ethanol and allow it to dry completely between each animal trial.
e. Sacrifice animals 90 minutes after behavior for c-Fos staining (if needed).

#### 12. Open-Field Test (OFT): Locomotion test

a. Place each animal individually in the center of the open-field arena (40 × 40 cm).
b. Record locomotor activity for 5 minutes using an overhead camera.
c. Quantify total distance traveled and average speed (e.g. ANY-maze).

#### 13. Elevated Plus Maze (EPM): Anxiety-like behavior test

a. Place each animal at the center of the EPM facing diagonal between open and closed arms.
b. Record behavior for 5 minutes.
c. Quantify time spent in the open arms.

#### 14. Y-maze: Short-Term Spatial Memory

a. Block one arm of the Y-maze (the designated novel arm) with a barrier.
b. Place the animal at the base of the maze and allow free exploration of the two accessible arms for 5 minutes (training phase).
c. Counterbalance which arm is blocked across animals and groups.
d. Return the animal to its home cage after training.
e. Clean the arena with 70% alcohol and wipe it dry.
f. After a 5-minute inter-trial interval, remove the barrier and reintroduce the animal to the Y-maze with all three arms accessible. The previously blocked arm is now the novel arm; the previously explored arm becomes the familiar arm.
g. Record behavior for 2 minutes (test phase).
h. Calculate spatial memory preference as the percentage of time spent in the novel arm relative to total time spent exploring both the novel and familiar arms combined:

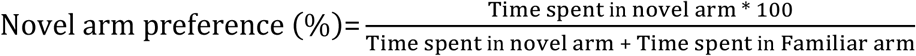

## Results

### Viral expression in targeted subpopulation of CA1 neurons

Confocal imaging of dorsal CA1 sections from CaMKIIα-CreERT2 mice (Fig. 5A) injected with the Flex-OFF hM4Di-mCherry virus showed mCherry fluorescence exclusively in neurons lacking EGFP signal. EGFP-positive neurons (Cre-expressing, CaMKIIα+) were devoid of mCherry signal (Fig. 5B, C), confirming successful Flex-OFF recombination and mutually exclusive subpopulation labeling at single-cell resolution. In contrast, injection of the complementary Flex-ON hM4Di-mCherry virus resulted in selective expression of mCherry in EGFP-positive neurons, confirming Cre-dependent targeting in dorsal CA1 (Fig. 5D, E).

**Figure 5.**
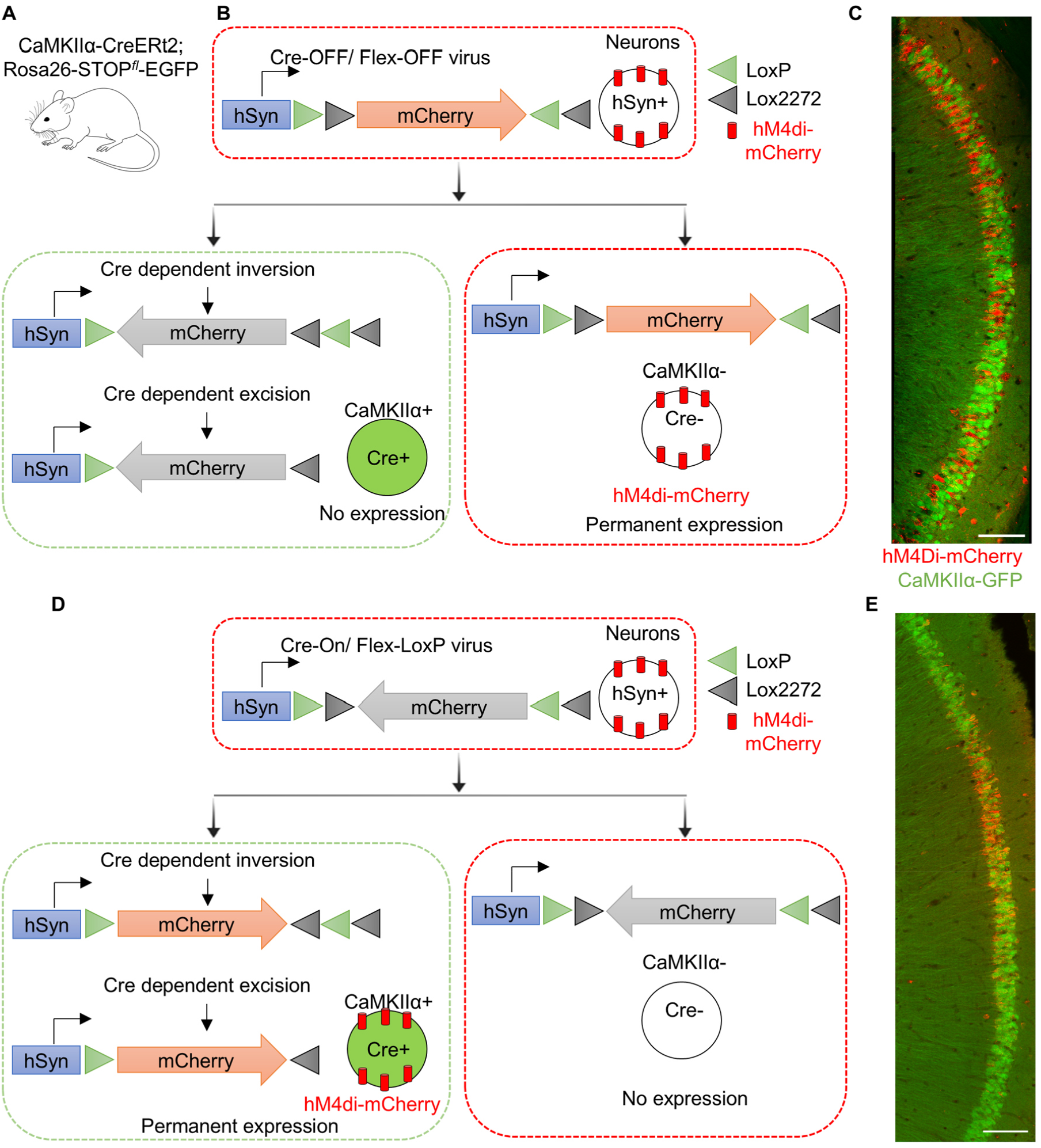
in-vivo Validation of Cre-Dependent AAV Expression. (A) Mouse-line used to genetically label excitatory neurons in the hippocampus using inducible Cre. (B) AAV (Flex-OFF) hM4Di-mCherry construct under the hSyn promoter is injected into CaMKIIα-CreERT2 mice. Schema showing viral expression in Cre+ neurons and Cre-neurons. (C) After Cre activation, hM4Di-mCherry expression was abolished in CaMKIIα+/Cre+ neurons and remained exclusively in CaMKIIα-/Cre-neurons. No colocalization of mCherry with GFP (marks Cre+ neurons) was observed, confirming Cre-off specificity. (D) AAV (Flex-ON) hM4Di-mCherry construct under the hSyn promoter is injected into CaMKIIα-CreERT2 mice. Schema showing viral expression in Cre+ neurons and Cre-neurons. (E) After Cre activation, hM4Di-mCherry was only expressed in Cre+ neurons, while Cre-neurons did not express. Colocalization of mCherry with GFP (marks Cre+ neurons) was observed, confirming Cre-ON specificity.

### Functional Validation of hM4Di-Mediated Inhibition in CA1 Neurons

Whole-cell patch-clamp recordings from hM4Di-mCherry-positive (Fig. 6C) CA1 pyramidal neurons showed a stable resting membrane potential prior to CNO application. Consistent with the previously published validation literature^30,31^, following bath application of CNO (10 μM), a hyperpolarization of the resting membrane potential was observed after 10 minutes of drug application, reflecting Gi-mediated activation of inwardly rectifying potassium channels (GIRKs) downstream of hM4Di receptor engagement (Fig. 6D). We also confirmed attenuation of action potential at the 125 pA current injection step after CNO application compared to pre-drug baseline recordings (Fig. 6E). This confirms functional expression of the hM4Di-mCherry construct and validate the capacity of the expressed receptor to suppress neuronal excitability in virally-targeted CA1 neurons.

**Figure 6.**
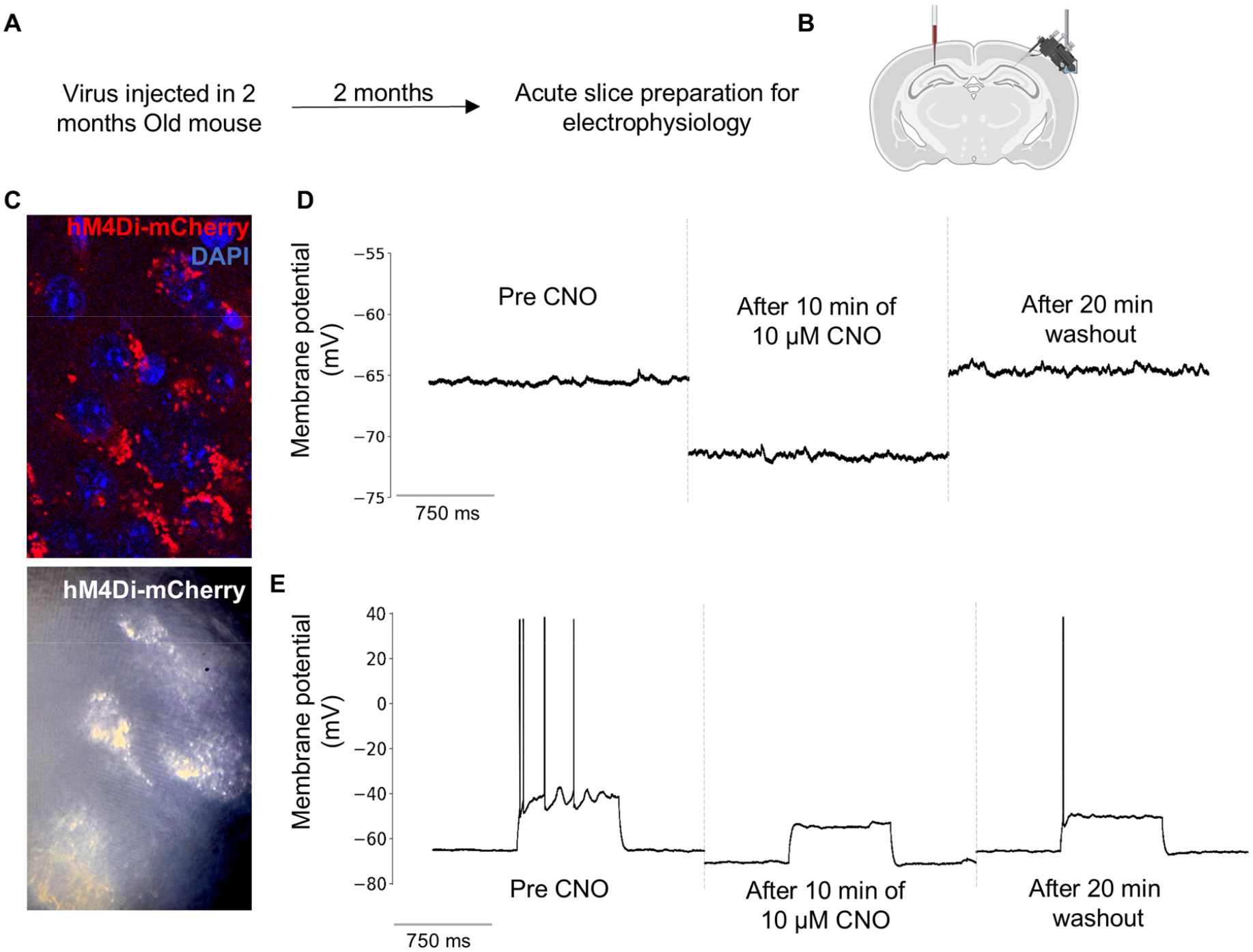
In-vivo functional validation of Cre-dependent hM4Di-mCherry expression. (A) Timeline of experimental design: AAV was injected into 2-month-old mice, and acute brain slices were prepared after 2 months for electrophysiological recordings. (B) Schematic of acute hippocampal slice showing the site of injection and patch-clamp recording. (C) Representative confocal image showing hM4Di-mCherry expression in hippocampal neurons, counterstained with DAPI (upper panel), and DIC image showing hM4Di-mCherry as bright spots (lower panel). (D) Whole-cell patch-clamp recording from an hM4Di-mCherry^+^ neuron shows hyperpolarization of the resting membrane potential after 10 min application of 10 μM CNO, with recovery after 20 min washout. (E) Current-clamp recordings from hM4Di-mCherry^+^ neurons (DIC image shown, left) demonstrate suppression of action potential firing after 10 min CNO application, with partial recovery after 20 min washout.

### CNO dose titration revealed locomotor suppression at higher doses

Prior to cognitive testing, we performed CNO dose titration test to rule out any potential locomotor side-effects, as higher doses of CNO have been shown to impair locomotion and produce off-target effects^32,33^. Virally injected animals are not required for this assessment. Animals received intraperitoneal injections of CNO at doses of 1, 5, or 10 mg/kg, 30 minutes prior to test locomotion using OFT. Total distance travelled and average speed was calculated using ANY-maze software.

Both the 5 and 10 mg/kg doses significantly reduced locomotor activity, resulting in decrease total distance travelled and reduced average speed, whereas the 1 mg/kg dose had no effect on locomotor activity (Figure 7A-C). Based on these findings, 1 mg/kg CNO was selected for use in all subsequent experiments.

**Figure 7.**
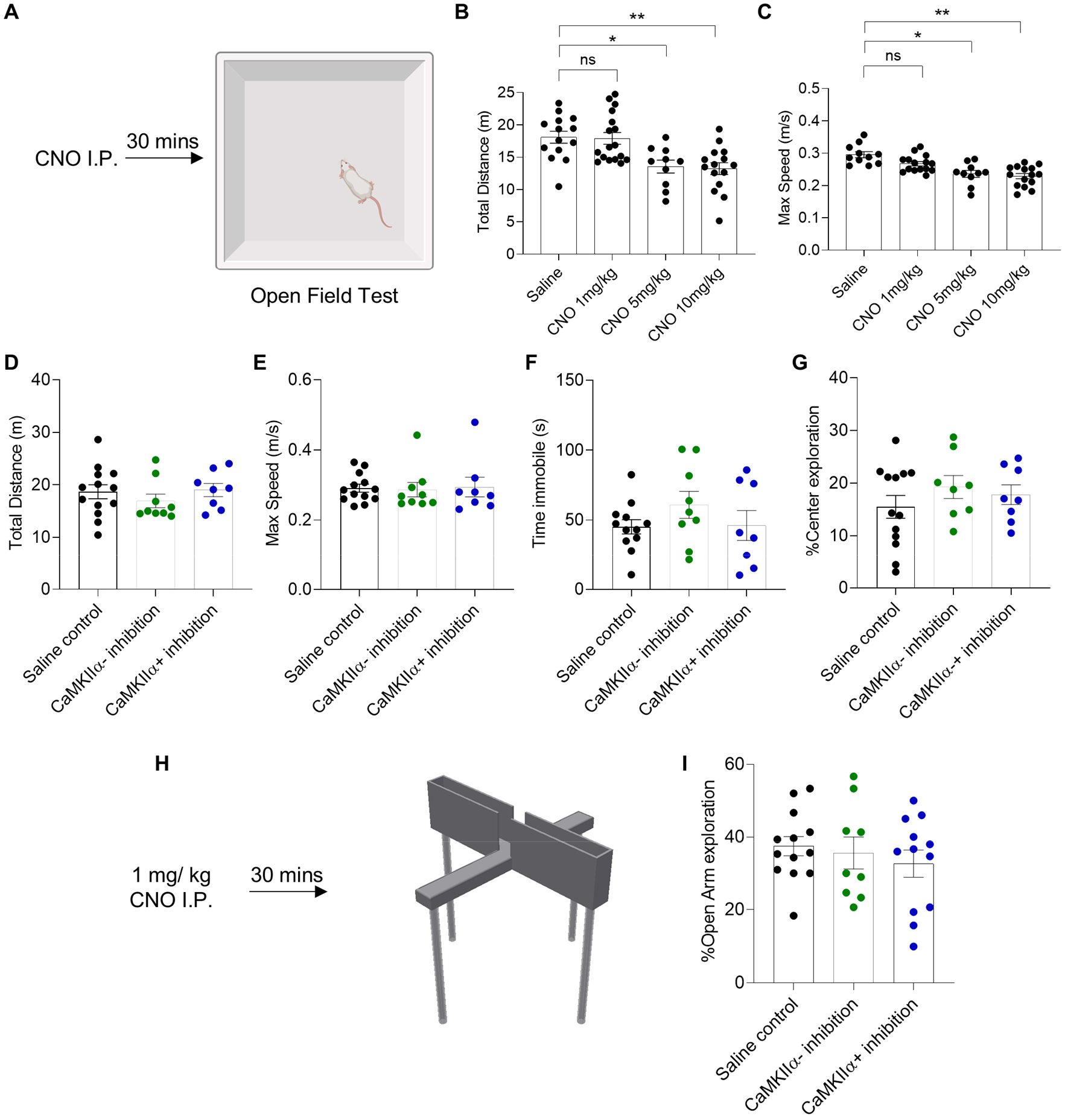
No effect of CaMKIIα+ or CaMKIIα-subpopulation inhibition on locomotion, and anxiety-like behavior. (A) Schema showing the time of CNO injection before OFT. CNO was injected at a dose of 1, 5, 10 mg/kg. (B-C) CNO dose titration shows both 5 and 10 mg/kg dose impairs total distance travel (B), and max speed (C), while 1 mg/kg dose did not cause any impairment in locomotion. N = 10-17 animals per group. 1 mg/kg was used in subsequent experiments. (D-E) Inhibition of either CaMKIIα+ or CaMKIIα-neurons in the dorsal CA1 does not impair mobility. No significant different was found in total distance travelled (D), and max speed (E) in OFT. (F-G) Neither CaMKIIα+ or CaMKIIα-inhibition impaired baseline anxiety. No difference was seen in time immobile (F), and center exploration (G) in OFT. N = 8-13 animals per group. (H-I) CaMKIIα+ or CaMKIIα-inhibition does not affect anxiety-like behavior in EPM. No differences were observed in open arm exploration in any group. N = 9-13 animals per group. Each dot represents an animal, and error bar represent mean ± SEM. One-way ANOVA, *p < 0.05, **p < 0.01, and ns is non-significant.

### Inhibition of CaMKIIα+ or CaMKIIα-neurons does not alter locomotion or anxiety-like behavior

Chemogenetic inhibition of either CaMKIIα+ or CaMKIIα-dorsal CA1 subpopulations did not alter baseline locomotor activity in the OFT. No significant differences were observed in total distance traveled, maximum speed, immobility time, or center exploration between saline and CNO treated groups (Fig. 7D-G).

Similarly, inhibition of either neuronal population did not affect anxiety-like behavior in the elevated plus maze. No differences were observed in open-arm exploration between groups (Fig. 7H, I), indicating neither CaMKIIα+ or CaMKIIα-alters anxiety-like behavior.

### CA1 subpopulation manipulation show differential impact on spatial memory

Previously published literature has shown that inhibition of CaMKIIα-positive neurons in dorsal CA1 disrupt spatial learning and memory^34,35^. However, how inhibition of CaMKIIα-negative neurons will affect short-term spatial learning and memory is not known. Animals were tested in the Y-maze novel arm exploration task following chemogenetic inhibition to test short-term spatial memory (Fig. 8A).

**Figure 8.**
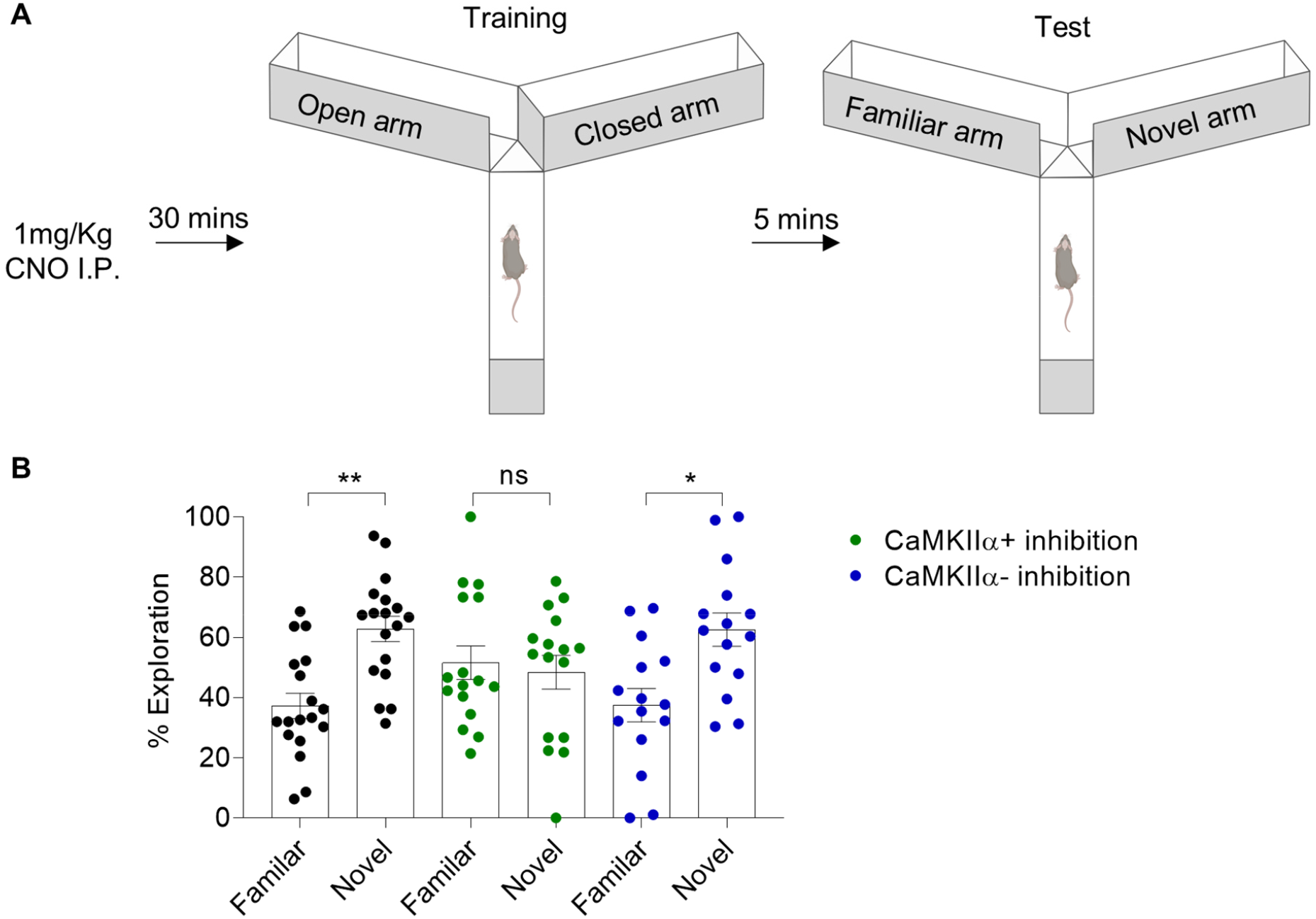
CaMKIIα+ inhibition in dorsal CA1 impair short term spatial learning and memory, while CaMKIIα-inhibition had no effect. (A) Schema showing the time of CNO injection before the training phase of Y maze. Animals were tested for spatial memory after a gap of 5 minutes. (B-C) Inhibition of CaMKIIα+ neurons caused an impairment in the short-term spatial learning or memory; no significant preference was seen for the novel arm. Both saline and CaMKIIα-animals showed a preference to novel arm implying normal spatial learning or memory. N = 15-18 animals per group. Each dot represents an animal, and error bar represent mean ± SEM. Paired t-test, *p < 0.05, **p < 0.01, and ns is non-significant.

Saline-treated animals showed a significant preference for the novel arm over the familiar arm, indicating intact short-term spatial memory. Inhibition of CaMKIIα+ neurons abolished this preference, demonstrating impaired spatial memory performance. In contrast, inhibition of CaMKIIα− neurons did not impair novel arm exploration, and these animals retained a significant preference for the novel arm comparable to saline-treated controls (Fig. 8B).

### Limitations

a. The in-house calcium-phosphate transfection method is sensitive to reagent quality and pH. Titer yields may be lower compared to commercial kit-based methods, larger-scale production for studies requiring very high viral doses.
b. The sucrose cushion ultracentrifugation protocol provides adequate purity for in vivo injection but does not achieve the same level of purity as iodixanol gradient ultracentrifugation. Residual cellular debris may cause elevated immune responses at the injection site in sensitive mouse strains.
c. The Flex-Off strategy is dependent on complete and irreversible Cre-mediated recombination. Incomplete tamoxifen induction in CaMKIIα-CreERT2 mice may lead to partial recombination, resulting in mosaic expression where some Cre-positive neurons still express hM4Di-mCherry. Adequate tamoxifen dosing and a sufficient waiting period post-induction are critical.

## Troubleshooting

### Problem 1: Low or absent fluorescence following transfection

#### Potential solution

- Confirm mCherry expression at 48 hours.
- Verify that the 2X HBS pH and ensure CaCl_2_ solution is freshly prepared (within 1 week) and stored at 4°C.
- Confirm cell confluency was approximately 70% at time of transfection. Over-confluent or under-confluent cells transfect poorly.
- Check that cells are within passage 10 and were passaged at least twice after revival from cryostock.
- Perform a positive control transfection using a known-working plasmid encoding GFP.

### Problem 2: Low viral titer after purification

#### Potential solution

- Increase number of T25 flasks per production batch; scale production proportionally.
- Ensure media collection is performed at all three time points (48, 72, and optionally 96 hours). Harvesting only at one time point substantially reduces yield.
- Verify that the sucrose cushion was properly formed before ultracentrifugation. If mixing occurred, repeat with fresh sucrose solution and careful technique.
- Increase centrifugation time to up to 4 hours.

### Problem 3: Off-target injection or poor viral spread

#### Potential solution

- Re-calibrate skull leveling before each injection session. Head tilt errors of even 0.1 mm in the anterior-posterior plane can shift targeting by >0.3 mm in deep structures.
- Perform dye injection calibration as described in the calibration step to confirm coordinates for your specific mouse strain and age.
- Reduce injection volume or rate if excessive spread is observed. Increase volume if labeling density is insufficient.
- Ensure the capillary tip is not blocked; cut tip or replace if consistent air pulses fail to produce a visible droplet.

### Problem 4: Incomplete Cre-mediated recombination / mCherry expression in EGFP+ neurons

#### Potential solution

- Verify tamoxifen induction protocol for CaMKIIα-CreERT2 mice. Standard induction requires 5 mg/ 35g tamoxifen I.P. for 7 consecutive days; allow at least 2 weeks after induction.

### Problem 5: No hyperpolarization following CNO application in electrophysiology

#### Potential solution

- Confirm that CNO stock solution is prepared correctly (10 mM in Saline or PBS) and that working concentration in aCSF is 10 μM. Check the expiry date and storage conditions of CNO.
- Allow the CNO bath application to continue for the full 10 minutes at 3-5 ml/min perfusion rate.

### Problem 6: Animals fail to show novel arm preference in Y-maze

#### Potential solution

- Test animals at consistent times of day.
- Verify that the arena is cleaned thoroughly between trials and that the testing room has consistent lighting and minimal noise.
- Use sufficiently distinct visual spatial cues in the testing room; absence of reliable extra-maze cues impairs spatial learning.

## Author contributions

DS standardized all methods and conducted all experiments described in the manuscript. DS wrote the manuscript. HSG supervised the project, edited manuscript, and provided funding. The authors have no competing interest to declare.

